# STPath: A Generative Foundation Model for Integrating Spatial Transcriptomics and Whole Slide Images

**DOI:** 10.1101/2025.04.19.649665

**Authors:** Tinglin Huang, Tianyu Liu, Mehrtash Babadi, Rex Ying, Wengong Jin

**Author notes:** Contributing authors.

## Abstract

Spatial transcriptomics (ST) has shown remarkable promise in pathology applications, shedding light on the spatial organization of gene expression and its relationship to the tumor microenvironment. However, its clinical adoption remains constrained due to the limited scalability of current sequencing technologies. While recent methods attempt to infer ST from whole slide images (WSIs) using pretrained image encoders, they remain restricted by limited gene coverage, organ-specific training, and the need for dataset-specific fine-tuning. In light of this, we introduce **STPath**, a generative foundation model pretrained on a large-scale collection of WSIs paired with ST profiles. This extensive pretraining enables STPath to directly predict gene expression across 38,984 genes and 17 organs without requiring downstream fine-tuning. STPath integrates multiple data modalities, including histological images, gene expressions, organ type, and sequencing technology information, within a novel geometry-aware Transformer architecture. Unlike previous methods that directly map WSIs to gene expression, STPath is trained using a masked gene expression prediction objective guided by tailored noise schedules, effectively balancing between capturing gene-gene dependencies and performing high-quality predictions. We evaluate STPath across 6 tasks spanning 23 datasets and 14 biomarkers, including gene expression prediction, spot imputation, spatial clustering, biomarker prediction, gene mutation prediction, and survival prediction. These results demonstrate STPath’s strong ability to infer spatially resolved gene expression and reveal crucial pathological structures within tissue samples, underscoring its promise for scalable ST-based pathology applications.

## 1 Introduction

Advances in computational pathology have enabled numerous clinical applications [68, 16, 3], and its integration with spatial transcriptomics (ST) [71, 52, 19] shows great potential in characterizing the tumor microenvironment [58, 5, 25] by capturing spatially resolved gene expression changes within the morphological context. Such multi-modal tissue representations offer finer-grained insights into cell-cell interactions and significantly facilitate biomarker discovery [37, 76, 10, 30]. However, conventional ST technologies remain low-throughput, and this challenge is further compounded by the large size of digitized tissue sections (whole-slide images, WSIs), limiting the availability of ST applications [59, 44, 18].

In light of this, several machine learning approaches have been proposed to infer spatially resolved gene expression directly from WSIs [24, 72, 12, 39]. These approaches leverage pathology foundation models [9, 73, 40, 80] pretrained on large-scale digital pathology collections [64, 13] to extract the representation for each spot segmented from WSIs. A learnable prediction head is typically trained to map these representations to gene expression profiles on small, organ-specific datasets, each tied to a particular sequencing technology and gene panel. As a result, the resulting models are narrowly specialized and cannot generalize well across different biological contexts. Moreover, their reliance on dataset-specific fine-tuning further restricts scalability, as each new deployment requires retraining the model to match the target gene set or sequencing protocol. To advance the applicability of ST in pathology, a model that generalizes across ST contexts without requiring repeated downstream adaptation is necessary.

In this work, we propose STPath, a foundation model for inferring spatial gene expressions from WSIs through a generative modeling paradigm. To train STPath, we assembled a largescale corpus of spatial transcriptomics and histology images from two publicly available datasets, i.e., HEST-1K [32] and STImage-1k4m [8], comprising 983 slides, 38,984 genes, 17 organs and 4 sequencing technologies. Extensive pretraining on this corpus enables STPath to learn a unified, generalizable representation across diverse organs and genes, allowing it to accurately predict spatial gene expression without requiring dataset-specific fine-tuning.

STPath represents each spot by fusing visual features, gene expression, organ type, and sequencing technology, and encodes all spots within a whole slide using a frame averaging-based spatial Transformer architecture [48, 28]. The model is pretrained through a masked modeling objective that recovers gene expressions for masked spots, guided by three sampling strategies: (1) an uniform sampling for selecting slide regions, (2) a Beta distribution for masking spots, and (3) a balanced schedule for gene prediction, alternating between highly variable genes and the full gene set. This pretraining paradigm leads to representations for WSIs that effectively encode the correlations between gene expressions and tissue morphology.

We evaluate STPath on 6 downstream tasks: gene expression prediction, spot imputation, spatial clustering, biomarker prediction, survival prediction, and gene mutation prediction. Compared to state-of-the-art methods [9, 73, 12, 72], STPath achieves the highest Pearson correlation on HEST-Bench [32], with a 6.9% improvement over the next-best approach, and further boosts performance with limited prompting, e.g., it gains +0.257 with just 5% of prompted spots on the CCRCC dataset. STPath also enhances pathology foundation models in weakly supervised settings, improving survival prediction by 6.1% for UNI and 5.6% for Gigapath, and gene mutation prediction by 5.6% for UNI and 5.0% for Gigapath among 8 datasets and 9 biomarkers. Additionally, our results demonstrate that the predicted biomarker expressions indicate mutation status accurately without further fine-tuning (e.g., TP53 in breast cancer reaches AUC = 0.73). These results highlight the effectiveness of STPath as a foundation model for integrating spatial transcriptomics in pathology applications.

## 2 Results

### 2.1 A Foundation Model for Spatial Transcriptomics-Pathology

STPath is pretrained on whole-slide images with spatial transcriptomics annotations. The model is optimized to predict the masked gene expression, and its main application is to infer the spatially resolved gene expression of certain or all the spots based on the pathology images, and benefits the clinical applications with the inferred expression, as shown in Figure 1(b). Additionally, pretraining on large-scale WSI collections allows STPath to learn representations for extensive genes. The pretrained model weights are available in a public hub, allowing users to leverage STPath for a wide range of downstream applications.

**Figure 1:**
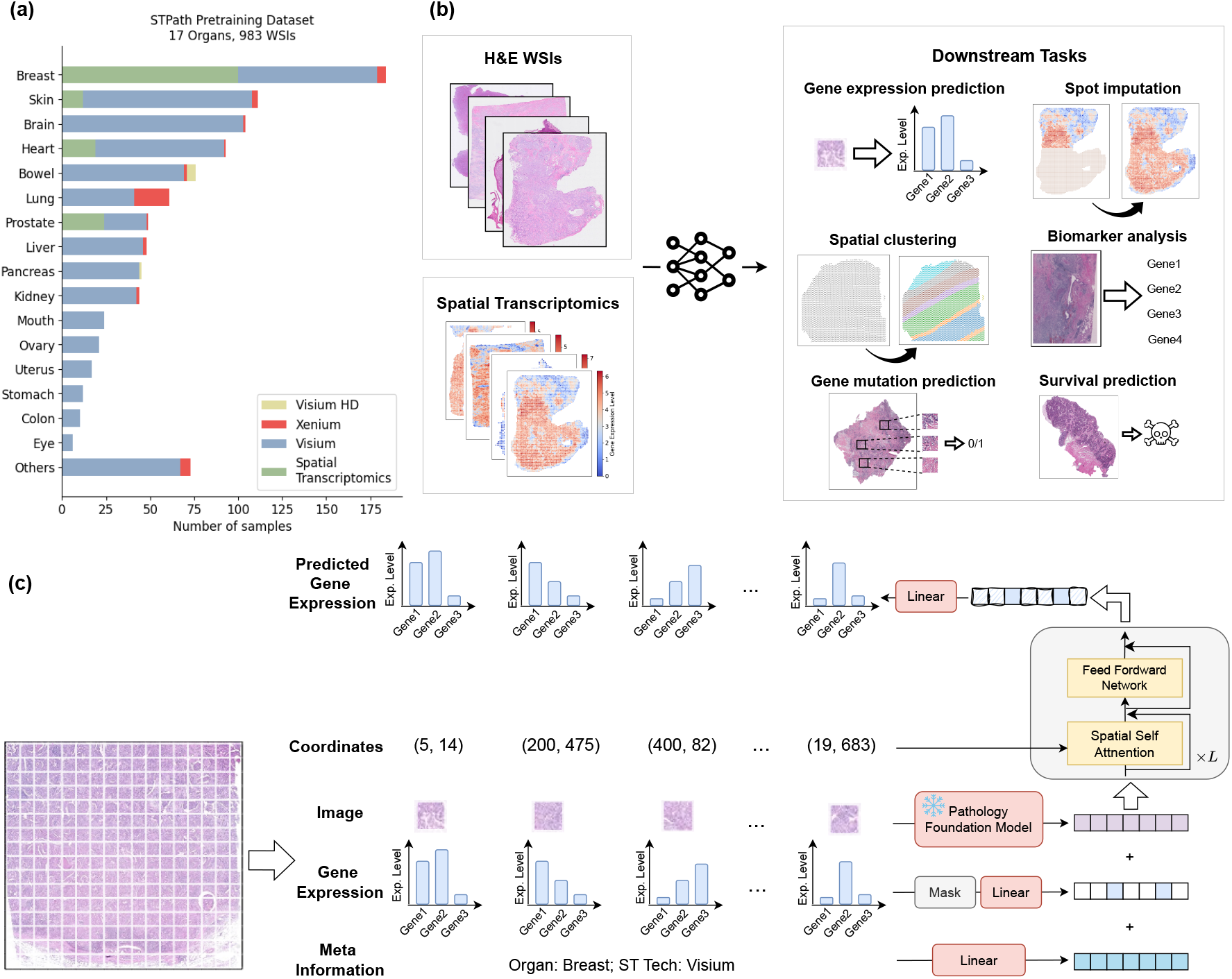
An overview of STPath. **(a)**: The statistics of datasets used for STPath pretraining, including the number of WSIs, the organ distributions, and the sequencing technologies. **(b)**: The six downstream applications of STPath, which incorporates the information of WSIs and the corresponding ST. **(c)**: The pretraining framework of STPath. A whole-slide image is first segmented into small spots, which will be encoded by a pathology foundation model. A portion of the gene expression data is masked, and the representations of all modalities are aggregated before being fed into the model. The spatial Transformer uses the coordinates to bias the attention map and infers the gene expression for the masked spots.

#### Dataset

We curated a large-scale dataset of paired H&E images and spatial transcriptomics annotations by integrating two public cohorts: HEST-1K[32] and STImage-1k4m[8]. To prevent label leakage and sample duplication, we removed samples in STImage-1k4m that originate from the same sources as HEST-1K. Additionally, samples that appear in downstream applications, e.g., HEST-Bench and spatial clustering, were excluded, resulting in 983 WSIs and 38,984 genes, as shown in Figure 1(a). We further set aside 5% of samples for validation, leaving 928 WSIs for pretraining. Additional details can be found in the Appendix A.2.

#### Methodology

STPath treats every spot of a WSI as a token, and all the associated modalities, i.e., image, gene expressions, organ type, and sequencing technology, are input as separate tracks that are aggregated into a representation. We apply the pathology foundation model, i.e., Gigapath [73], to embed the spot images. A spatial-aware Transformer architecture is employed as the model backbone, which biases the attention map with spatial dependency using an invariant frame averaging-based transformation [48, 28]. During training, some gene expressions are masked following a specific distribution, and the model is supervised to predict the expression levels of certain genes. The objective is a regression loss, which differs from the classical masked language modeling approach [35]. Further technical details can be found in Section 4.

### 2.2 STPath accurately infers spatially-resolved gene expressions

We evaluate the performance of STPath on the task of gene expression prediction and validate the effectiveness of its architecture by comparing it to several state-of-the-art baseline methods, including UNI [9], Gigapath [73], BLEEP [72], and TRIPLEX [12], which are trained on the same cohort for a fair comparison. We apply HEST-Bench [32] that includes 10 cancer datasets with different organs. As shown in Figure 2(a,b), rather than following the original leave-one-out cross-validation setup, we report the average performance among all the samples in each dataset without further finetuning to ensure a more realistic evaluation. The Pearson correlation coefficient (PCC) and adjusted mutual information (AMI) between the predicted and ground truth expression levels of the top 50 highly variable genes (HVGs) are used as evaluation metrics. Additional details regarding the experimental setup and implementation are provided in Appendix A.1 and Section 4.4, respectively.

**Figure 2:**
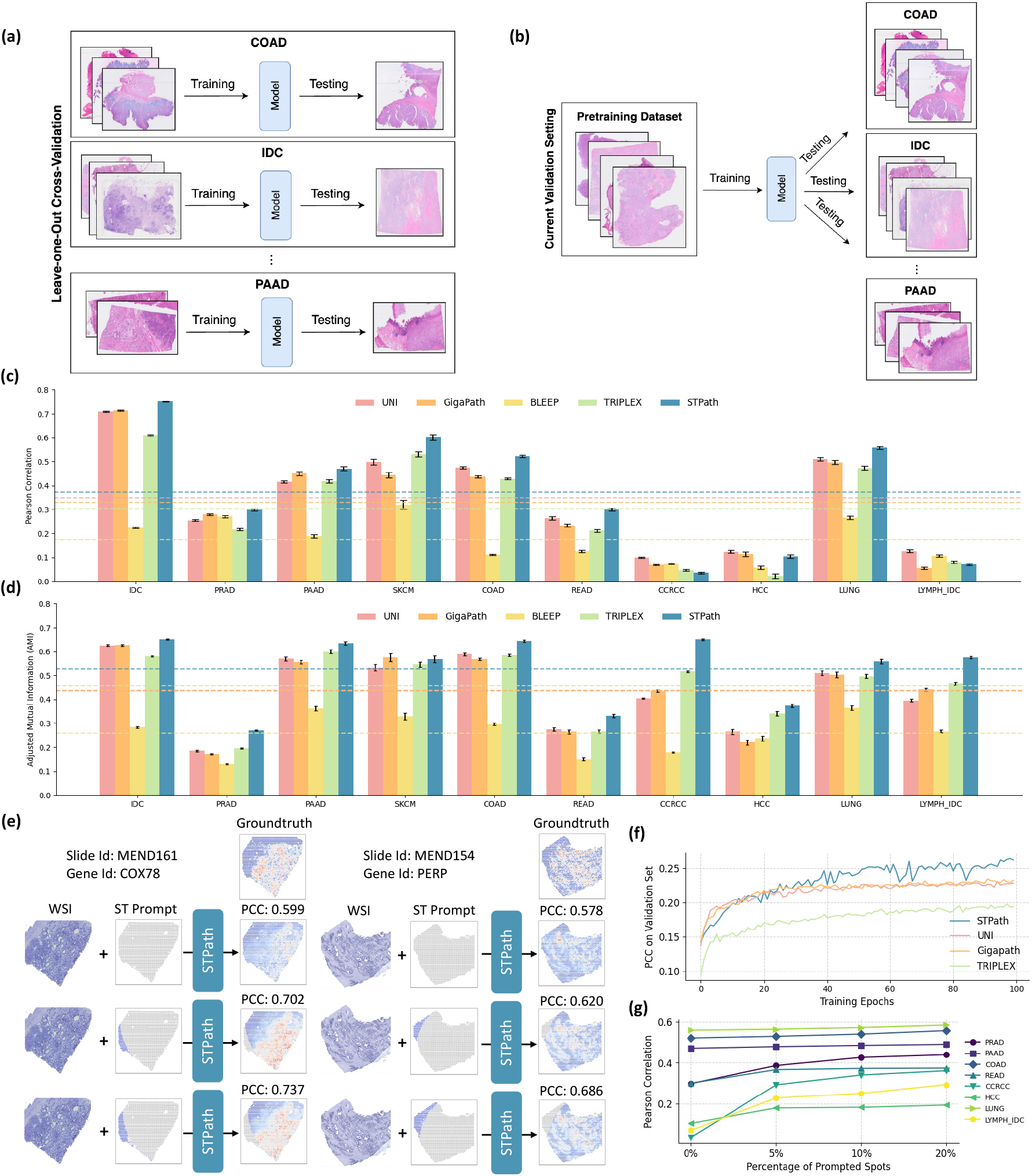
Gene expression prediction on HEST-Bench. **(a)**: The original evaluation setup used in HEST-Bench, where a separate model is trained for each fold of each dataset. **(b)**: The evaluation setup used in our work, where a single model is pretrained on a large-scale dataset and evaluated in a zero-shot manner. **(c**,**d)**: Performance on the top 50 highly variable genes measured by Pearson correlation (c) and adjusted mutual information (d). The dashed lines indicate the average performance of each model across all datasets. **(e)**: Two cases from HEST-Bench on how STPath receives prompted gene expressions and produces more accurate predictions. **(f)**: The average Pearson correlation of the top 200 highly variable genes on the validation set as pretraining steps increase. **(g)**: Performance of STPath with varying percentages of prompted spots.

During pretraining^1^ (Figure 2(f)), STPath outperforms all baseline models in terms of PCC on the top 200 HVGs from the validation set, with the performance gap widening over time. Compared to the slide-based model TRIPLEX (PCC = 0.198), STPath achieves a higher average PCC of 0.266, reflecting a +34.4% improvement. On comparison of HEST-Bench (Figure 2(c,d)), STPath achieves the highest average performance in both PCC (ranking first in 8 out of 10 datasets) and AMI (ranking first in 9 out of 10 datasets). On average, it outperforms the next best-performing models by 6.9% in PCC (UNI) and 14.4% in AMI (TRIPLEX), both with *P*-value *<* 0.001 evaluated by a two-sided Wilcoxon test. Notably, the PCC achieved by STPath is particularly strong on certain datasets, such as 0.75 on IDC, 0.60 on SKCM, and 0.559 on LUNG, highlighting its capability to accurately infer gene expressions across organs.

#### STPath effectively imputes gene expression with a small number of prompts

Unlike previous studies primarily focusing on learning a direct mapping from images to gene expressions, the generative learning paradigm allows in-context prediction, enabling the model to leverage the expressions of a few prompted spots for more accurate predictions. As illustrated in Figure 2(e), we assess the model’s performance in predicting the gene expressions of masked spots based on the expressions of the remaining visible spots. We select datasets where the model struggles to achieve significant correlation, i.e., PCC<0.7. The proportion of prompted spots varies within the range of [0%, 5%, 10%, 20%], with 0% representing the task of predicting expressions for all spots.

Across all datasets (Figure 2(g)), we observe a consistent improvement in PCC as the proportion of prompted spots increases. For instance, at just 5% prompting, CCRCC shows a notable absolute gain of +0.257, HCC improves by +72% (from 0.104 to 0.179), and PRAD increases by +29% (from 0.298 to 0.386). Moreover, a higher number of annotated spots (i.e., lower masking rates) consistently leads to better prediction accuracy. For example, in LYMPH_IDC, the PCC improves from 0.070 at 0% to 0.293 at 20%, representing a +318% increase. Similarly, READ sees an improvement from 0.300 to 0.374, and PAAD rises from 0.471 to 0.490 as prompting increases from 0% to 20%. The complete results can be found in Appendix A.3.

### 2.3 Gene expressions predicted by STPath capture cell type information

In this section, we evaluate the ability of STPath to infer and characterize spatial patterns within tissue samples, offering insights into disease prognosis. This study utilizes five datasets with pathologist annotations [42, 15, 8, 4, 20], resulting in a total of 42 samples. To quantify model performance, we employ metrics that measure the alignment between Leiden cluster assignments and annotations, including adjusted mutual information (AMI) and homogeneity score (Homo). The average performance across all the samples in one dataset is reported (Figure 3). The detailed experimental setup can be found in Appendix A.1.

**Figure 3:**
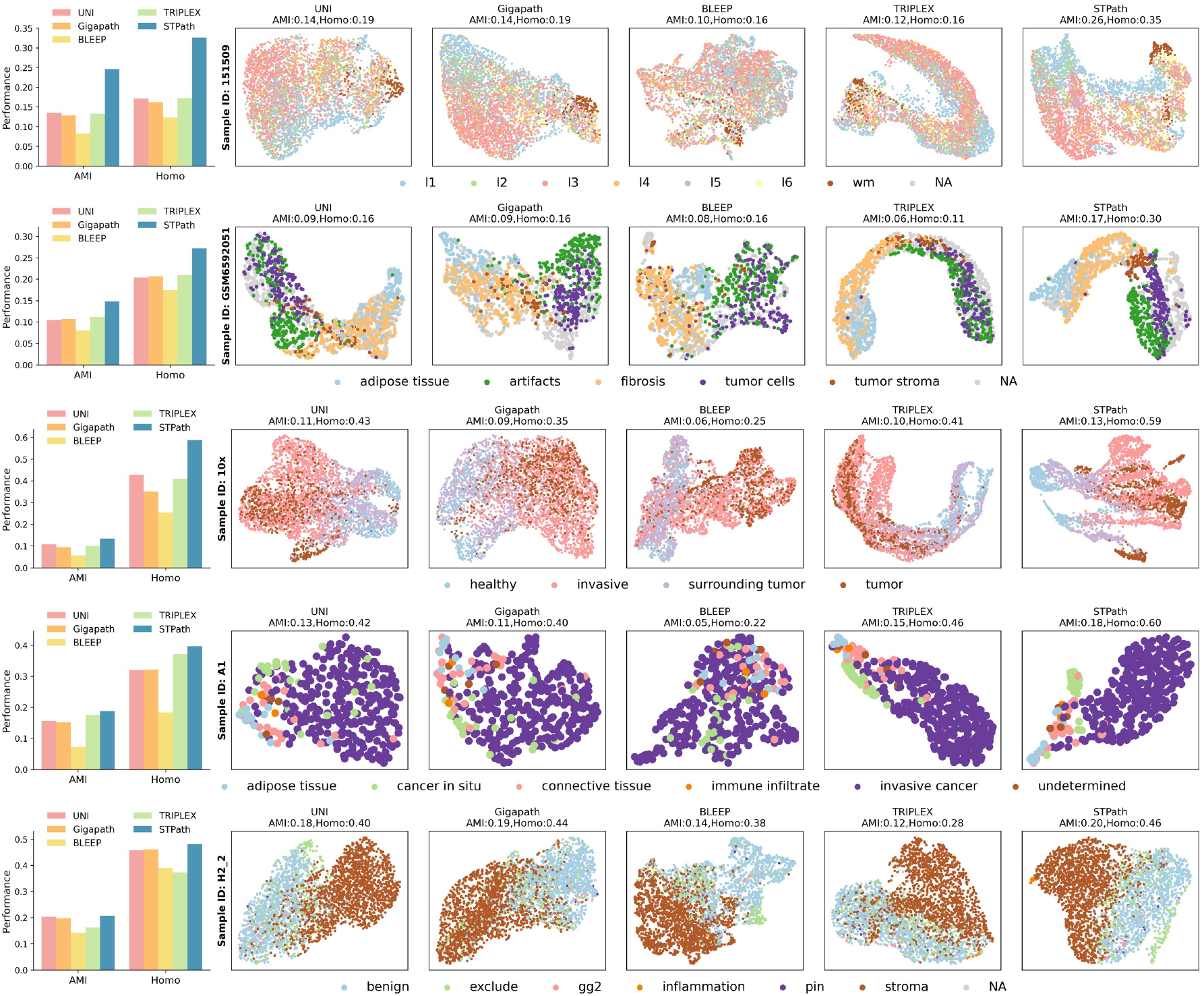
Clustering results on five datasets with pathologist annotations are shown: human brain samples with layer annotations [42], samples with breast cancer subtypes [15], samples with tumor annotations [8], samples with tumor cell types [4], and samples with benign and malignant tumor annotations [20]. One clustering example per dataset is presented in each row.

Compared to other methods, STPath consistently achieves the best performance across all datasets. For example, it outperforms the strongest baseline by a substantial margin of 0.121 on the human brain dataset and achieves a +29.6% improvement on the 10x sample. Qualitative visualizations further highlight that STPath produces more coherent and spatially aligned segmentations, effectively capturing biologically meaningful regions such as tumor cores, stromal compartments, and immune infiltration zones.

### 2.4 STPath accurately predicts cancer-related marker genes

The expression level of biomarkers serves as a critical indicator of tumor presence and progression. In Figure 4(a), we compare the Pearson correlation between the predicted and ground truth expressions for 6 known cancer biomarkers across 4 datasets from HEST-Bench, i.e., GATA3 and MYBPC1 on IDC [26, 11, 22], LCN2 on COAD [33], GPRC5A and SOX2 on LUNG [78, 55, 27], and UBE2C on PAAD [70, 41]. Compared to the other methods, STPath consistently achieves the best performance with +9.3% improvement over the next-best baselines (*P*-value *<* 0.001). Besides, STPath can achieve a promising correlation on GATA3 and MYBPC1 (PCC *>* 0.95), both well-established markers in breast cancer, highlighting the capability of STPath in capturing clinically meaningful gene expression patterns.

**Figure 4:**
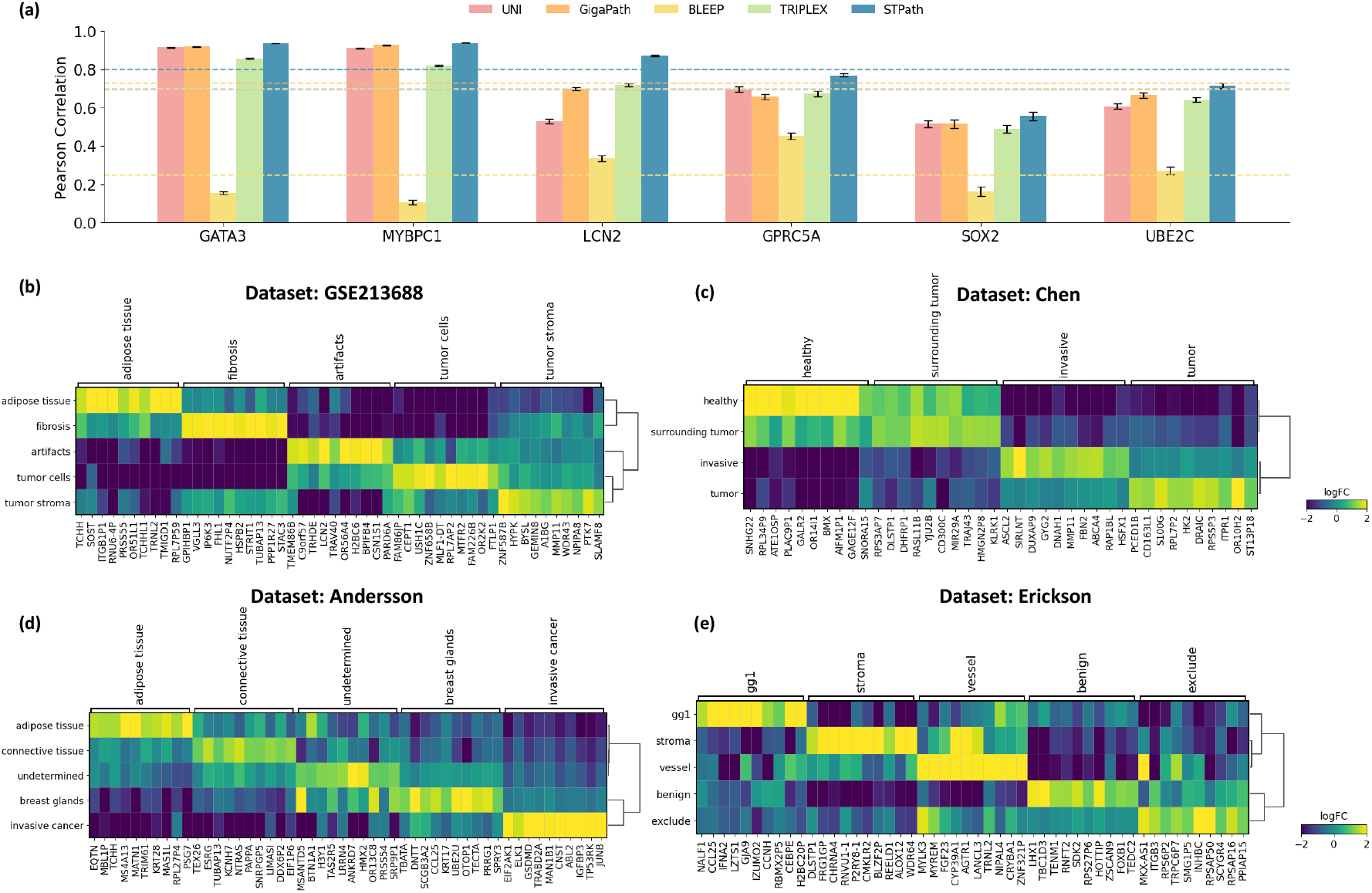
Experimental results for biomarker analysis. **(a)**: Comparison between STPath and baselines in predicting expressions among 6 biomarkers on 4 datasets from HEST-Bench. **(b-e)**: Matrix plots showing the log fold change (logFC) of the top 10 marker genes derived from the predicted spatial transcriptomics by STPath, aligned with pathologist annotations. Plots correspond to a representative sample from 4 datasets [42, 8, 4, 20].

Moreover, we assess whether STPath can identify tumor-relevant marker genes by performing Wilcoxon rank-sum tests on four representative cancer samples with pathologist annotations (Figure 4(b-e)). Our analysis reveals that STPath successfully recovers a range of known biomarkers. For instance, in the breast cancer samples (Figure 4(b-d)), several well-established tumor markers are highlighted, including SPAG5[21, 79], SLFN12[1], CDC7[60], OSCP1[29], MMP11[77, 45], S100G[53], GSDMD [38], and IGFBP3 [50]. In the prostate sample (Figure 4(e)), we observe enrichment of melanoma- or tumor-associated genes in malignant regions, such as CCL25 [2], IFNA2 [36], and CEBPE [74]. These findings demonstrate that STPath can uncover biologically meaningful markers aligned with known cancer signatures.

### 2.5 STPath enhances slide-level prediction via spatial transcriptomics

Predicting clinical outcomes from histology images, e.g., patient survival and gene mutation status, plays a critical role in computational pathology, offering insights into disease prognosis and enabling personalized treatment strategies [69, 57, 73]. However, these labels are typically assigned at the whole-slide level, rather than to the specific regions where relevant biological signals, such as biomarker mutations, are localized [9, 73].

In this section, we investigate the capability of spatially resolved gene expressions inferred by STPath to complement spatial context and enhance slide-based prediction. This study includes eight datasets: BRCA (invasive breast cancer) [7, 56], COAD (colon adenocarcinoma) [63], LUAD (lung adenocarcinoma) [46], GBM (glioblastoma multiforme) [14], LSCC (lung squamous cell carcinoma) [65], and HNSC (head-and-neck cancer) [43], MBC (metastatic breast cancer) [6], and SURGEN (Survival and Genetic Markers) [47], each split into training, validation, and test sets following a 6/2/2 patient-level split. The inferred expressions of 14 biomarkers are concatenated with visual features, and ABMIL [31] is employed as the prediction head (Figure 5(a)). More information regarding the experimental setup and datasets can be found in Appendix A.1 and Appendix A.2, respectively.

**Figure 5:**
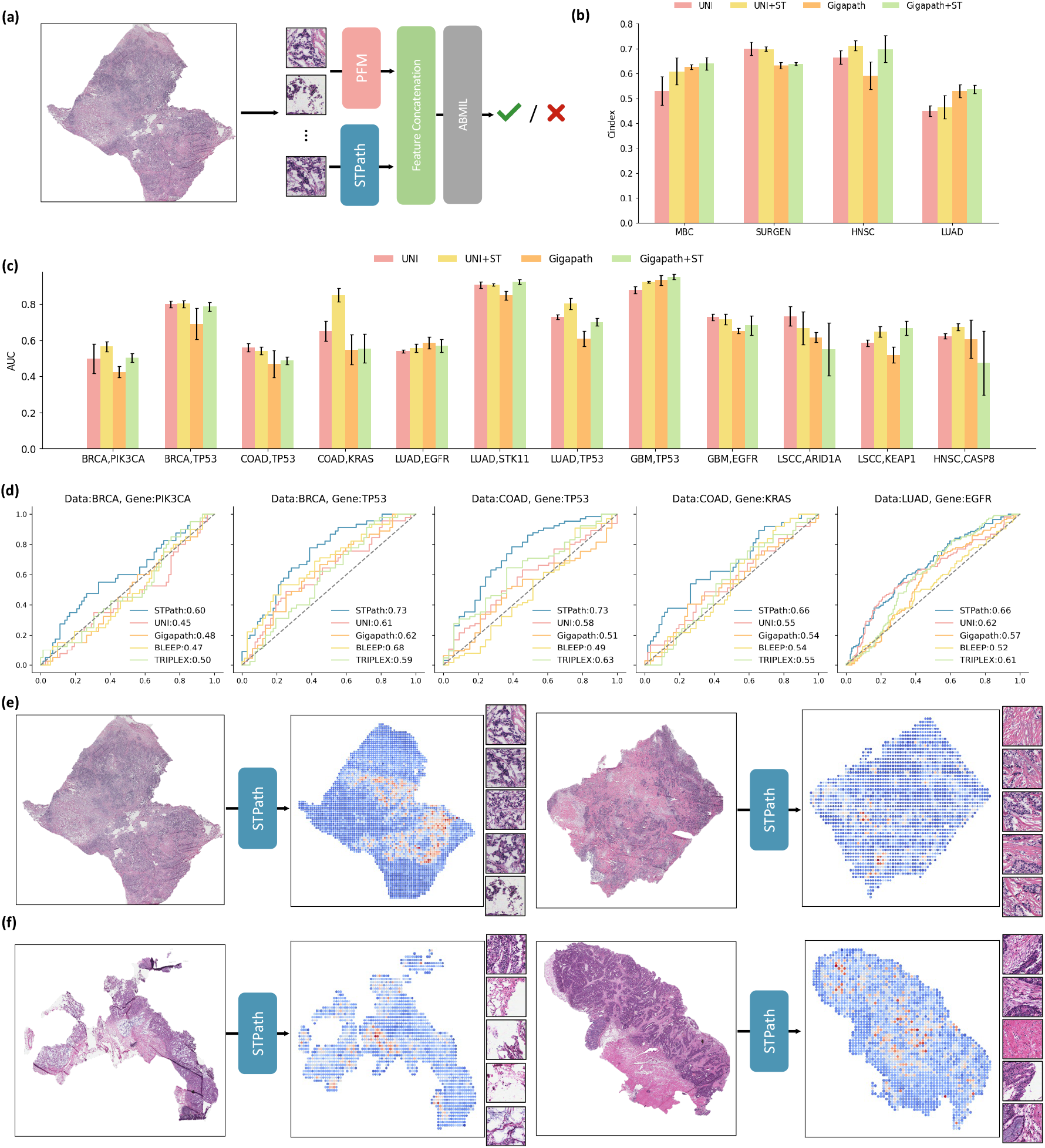
Results of survival prediction and gene mutation prediction. **(a)** Weakly supervised classification with STPath. Predicted spatial transcriptomics features by STPath are combined with visual features and passed to a multi-instance-learning head. **(b, c)** Performance of using predicted spatial transcriptomics as additional features in survival prediction (b) and gene mutation prediction (c) under the weakly supervised setting. **(d)** Comparison of AUC scores between STPath and baseline methods under the unsupervised mutation prediction setup. **(e, f)** Visual examples from BRCA (e) and COAD (f), showing the predicted expression of the biomarker and the corresponding patches with the highest expression levels.

#### Weakly Supervised Survival Prediction

Four datasets, including MBC, SURGEN, HNSC, and LUAD, are applied in this study, and the C-index is used as the evaluation metric. We observe consistent performance improvements in both pathology foundation models when incorporating the predicted spatial transcriptomics from STPath, with gains seen in 7 out of 8 cases. On average, this yields a 6.1% improvement for UNI and 5.6% for Gigapath (Figure 5(b)). The most notable improvement is observed on the HNSC dataset, where UNI increases from 0.665 to 0.771, and Gigapath from 0.591 to 0.698.

#### Weakly Supervised Gene Mutation Prediction

In addition to survival prediction, we also assess STPath on the gene mutation prediction task [49, 34, 51]. Six public datasets from Clinical Proteomic Tumor Analysis Consortium (CPTAC) [17, 61] for pan-cancer mutation prediction are applied: BRCA, COAD, LUAD, GBM, LSCC, and HNSC These datasets cover a range of biomarkers: TP53, KRAS, PIK3CA, EGFR, STK11, ARID1A, KEAP1, and CASP8. The area under the curve (AUC) is used as the evaluation metric. As shown in Figure 5(c), incorporating predicted gene expressions improves performance in the majority of cases, with gains observed in 18 out of 24 settings. On average, the predicted spatial transcriptomics from STPath leads to a performance increase of 5.6% for UNI and 5.0% for Gigapath.

#### Unsupervised Gene Mutation Prediction

Benefiting from the extensive pretraining, STPath can be applied to directly infer the expression of the studied biomarker for each spot. We interpret the variance in biomarker expression across spots within a WSI as a mutation indicator, which is then evaluated across all samples in each dataset. All the slides in each dataset will be used for evaluation without additional splitting.

We observe that the proposed unsupervised paradigm achieves promising results on five biomarker cases (Figure 5(d)): PIK3CA (AUC = 0.60) and TP53 (AUC = 0.73) on BRCA; TP53 (AUC = 0.73) and KRAS (AUC = 0.66) on COAD; and EGFR (AUC = 0.66) on LUAD. On average, it outperforms the best-performing baselines with a gain of +13.9%. Additionally, the spatially resolved gene expression inferred by STPath can highlight regions of high diagnostic importance. As shown in Figure 5(e,f), we present four examples from BRCA and COAD, displaying the predicted biomarker expression along with spot images exhibiting the highest expression levels.

## 3 Discussion

In this study, we propose STPath, a foundation model for integrating spatial transcriptomics with H&E-stained histology images. Unlike previous studies focusing on organ- or gene-specific cohorts, STPath aims to learn a universal representation capable of generalizing across diverse organs, genes, and sequencing technologies. Such a model is made possible through a generative pretraining paradigm over a large-scale collection of WSI-ST pairs. Specifically, we curated a largescale dataset by combining two public resources: HEST-1K and STImage-1k4m, each consisting of WSIs annotated with spatial transcriptomics. We model different data modalities, including spot images, gene expressions, organ types, and sequencing technologies, in separate tracks, and fuse them into unified spot representations. These representations are then processed by our proposed geometry-aware Transformer with spatial coordinates. A generative objective with dedicated masking schedules is employed to pretrain the model. Importantly, STPath is the first model demonstrating the effectiveness of the foundation model pretraining with WSI-ST pairs, enabling strong zero-shot and few-shot performance across a variety of downstream tasks and datasets.

We demonstrate the capability of STPath across multiple applications in spatial transcriptomics and clinical pathology. For gene expression prediction, STPath accurately infers gene expressions across 10 datasets of different organs, achieving strong correlations with ground truth values without additional fine-tuning. Moreover, the generative modeling paradigm enables improved prediction performance when prompted spots are provided, demonstrating its ability for spatial context understanding. The gene expressions predicted by STPath also exhibit strong alignment with pathological structures, achieving the highest correlation with expert annotations, such as brain layers and tumor regions, across five datasets. Furthermore, we apply the predicted biomarker expressions from STPath to determine mutation status across three cohorts, demonstrating promising accuracy relative to annotated labels. For instance, STPath achieves an AUC of 0.73 for TP53 mutation prediction in breast cancer samples.

Regarding the methodology, STPath offers several advantages over previous related studies: (1) it learns an extensive gene vocabulary comprising 38,984 genes, covering a broad spectrum of biomarkers and enabling strong generalizability across diverse gene panels and datasets; (2) it employs a geometry-aware Transformer architecture that is *E*(2)-invariant, allowing more effective encoding of WSIs at varying scales; (3) it demonstrates promising potential for improving clinical applications, including mutation and survival prediction, by providing spatially resolved gene expression as additional contextual information.

In conclusion, applying generative modeling and multi-modal pretraining to spatial transcriptomics prediction from H&E-stained images represents a promising yet largely underexplored direction. Recent advances in large-scale WSI-ST collections create a valuable opportunity to develop generative foundation models that bridge the gap between spatial transcriptomics and histopathology, motivating our proposal of STPath. While STPath shows strong performance across many datasets, it currently struggles to predict gene expression with high correlation for certain organs, such as CCRCC in HEST-Bench. We attribute this limitation to the scarcity of relevant data in the pretraining set. We expect to improve STPath s performance by incorporating more diverse datasets and additional modalities, such as scRNA-seq.

## 4 Methods

**Notation** The target of STPath is to infer the spatial transcriptomics of a given WSI. Specifically, an H&E-stained WSI is segmented into spots, which can be represented as *x* = (***x***_coord_, ***x***_gene_, ***x***_img_), with coordinates ***x***_coord_ ∈ ℝ^*N* ×2^, spot images ***x***_img_ ∈ ℝ^*N* ×3×*H*×*W*^, and gene expression levels ***x***_gene_ ∈ ℝ^*N* ×*G*^, where *N* is the number of spots, *G* is the total number of gene symbols, and *H, W* indicate the image dimensions.

### 4.1 Model Input

STPath represents each spot as a discrete token, and the token representation is obtained by aggregating its visual feature, gene representation, and meta information. Each modality is embedded as a separate track with distant learnable parameters.

#### Visual Feature

We employ the pathology foundation model Gigapath [73] to encode and extract the visual features from each spot. Gigapath consists of a patch-level encoder and a slide-level encoder; in this track, we use the patch-level encoder to embed the spot images, and an additional linear transformation is applied to align the latent space with the aligned dimension:

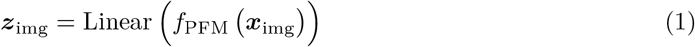

***z***_img_ = Linear where ***z***_img_ ∈ ℝ^*N* ×*d*^ and *d* is the hidden size.

#### Gene Representation

We cleaned and constructed a vocabulary consisting of 38,984 gene symbols, with each gene symbol assigned a unique ID. Gene expression can then be represented as a multi-hot vector using the constructed vocabulary, where the expression level of each gene is placed in its corresponding ID position. The gene expression is normalized by log1p transformation. We apply a linear transformation to aggregate the gene expressions into a representation:

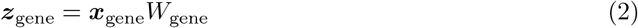

where ***z***_gene_ ∈ ℝ^*N* ×*d*^ and *W*_gene_ is a learnable transformation matrix. Given that gene expression is represented as a multi-hot vector, this linear transformation is equivalent to a weighted summarization over a bag of embeddings, where the weights correspond to the normalized expression values.

#### Meta Information

We incorporate sequencing technology and organ information into the modeling process. Specifically, each organ and sequencing technology is assigned a unique ID and replaced with a learnable embedding:

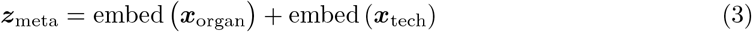

where ***z***_meta_ ∈ ℝ^*N* ×*d*^.

#### Spot Representation

The embeddings over all the tracks are summed into one representation as input to the first layer of the model architecture:

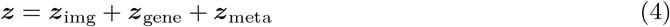

where ***z*** ∈ ℝ ^*N* ×*d*^ denotes the spot representations.

### 4.2 Geometry-Aware Transformer Architecture

STPath receives a sequence of spot representations along with their corresponding coordinates in WSI as input, which will be processed within a geometry-aware Transformer architecture. The proposed architecture employs an *E*(2)-invariant self-attention mechanism, ensuring invariance to rotations, reflections, and translations operated on the coordinates. This invariance is achieved through the frame averaging framework [48, 28]. Given a set of coordinates ***x***_coord_ ∈ ℝ ^*N* ×2^, we first compute the relative direction between all spot pairs:

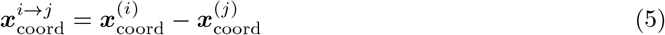

where 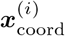 denotes the *i*-th spot’s coordinate. To simplify notation, we define the set of all pairwise direction vectors as 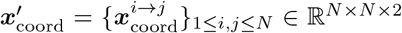 We further project this geometric context into four frames using PCA:

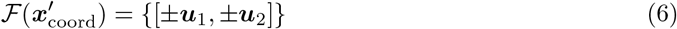

where ***u***_1_, ***u***_2_ are the two principle components extract from the given 2D point cloud. Building on top of the extract frames, we can perform an *E*(2)-invariant transformation by averaging across the frames:

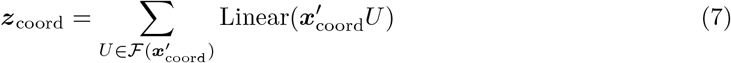

where ***z***_coord_ ∈ ℝ ^*N* ×*N*^ represents the projected scalar features that encode geometric information. These features are integrated into the self-attention mechanism as a spatial bias:

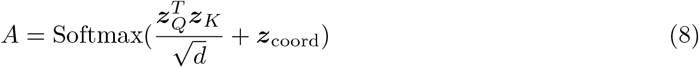

where ***z***_*Q*_, ***z***_*K*_ are the query and key representations projected from ***z***, and *A* ∈ ℝ ^*N* ×*N*^ is the calculated attention map. The spatial bias ***z***_coord_ is derived based on the relative direction vectors between spots, therefore encoding the distance and the orientation information. Similar to conventional Transformer [35, 67], the spot representations are aggregated using the calculated attention map and processed with Feed-Forward Network.

### 4.3 Pretraining Strategy

In the generative pretraining paradigm, corruption is applied to the part of gene expressions and the optimization objective of the model is to recover the masked gene expressions. MSE loss is applied as the loss function. The overall learning objective is shown below:

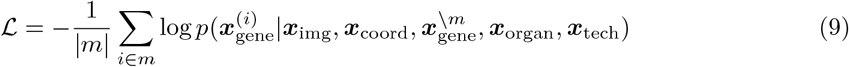

where *m* denotes the random mask applied to the gene symbols of slide. To ensure an effective and robust training process, we apply three different schedules for different phases of the model training.

#### Slide Sampling Schedule

To balance different scales of WSIs while maintaining efficiency, a continuous region is sampled from the slide at each training step and used as the model input. The number of spots is drawn from a Uniform(64, 1024) distribution. The sampling on slides can also be considered as a form of data augmentation, introducing variability and improving robustness. In our experiments, we found that training with relatively small regions enables the generalization to large slides. We attribute this to the models ability to learn localized spatial patterns effectively, which can then be extrapolated to broader contexts when processing entire WSIs.

#### Masked Noise Schedule

The goal of designing a masked schedule is to balance the utilization of gene-gene dependencies while ensuring high-quality predictions based on WSIs. A small masking ratio, such as the 15% used in BERT [35, 23], reduces the objective to spot imputation, where unmasked gene expressions serve as strong indicators, limiting the models ability to capture deeper dependencies. On the other hand, masking all spots prevents the model from leveraging gene-gene relationships.

To achieve a good trade-off, we sample the masking rate from a Beta(10, 1) distribution, ensuring that most spots are masked most of the time (approximately 90% on expectation) while still allowing for an exposure of gene expressions as input, mimicking a prompted input structure. For the masked spots, 95% are replaced with [MASK] token with a learnable embedding, while the remaining 5% have 50% of their gene expressions removed. Additionally, to ensure that STPath can perform well even when sequencing technology information is unavailable, we randomly replace the technology embeddings of spots with a learnable padding embedding with an 80% probability.

#### Gene Target Schedule

At each training step, we predict the expression level of all gene symbols over all spots but only calculate the loss among the genes that are measured in the sample. To minimize the impact of housekeeping genes [62, 54] on loss sensitivity, 80% of the time the model predicts the expression of highly variable genes, while the remaining 20% involves predicting the expression of all measured genes.

### 4.4 Implementation

The experiments are conducted on a single Linux server with the AMD EPYC 9534 64-Core Processor, 1024G RAM, and 8 NVIDIA L40S-48GB. Our method is implemented on PyTorch 2.3.1 and Python 3.10.16.

#### STPath

We use the Adam optimizer with a learning rate of 5×10^−4^ and mean squared error (MSE) as the loss function. To ensure learning stability, the gradient norm is clipped to 1.0 at each training step. The model is trained with early stopping, which is triggered if no performance improvement is observed on the validation set for 20 epochs. The model checkpoint with the lowest validation loss is selected as the final model. Each WSI is treated as a single training instance, with a fixed batch size of 2. The model architecture consists of 4 layers, a hidden size of 512, and 4 attention heads. Additionally, dropout is set to 0.1, and the activation function used is GeLU. Gigapath is applied as the histology image encoder since it generally exhibits better performance in our experiments. STPath is available at GitHub:https://github.com/Graph-and-Geometric-Learning/STPath.

#### Baselines

We follow the official implementation for each baseline: UNI^2^, Gigapath^3^, BLEEP^4^, and TRIPLEX^5^. To ensure a fair comparison, all models are trained using the same setup as STPath, including the loss function, optimizer, early stopping criterion, slide sampling strategy, and histology image encoder (Gigapath is used as the encoder for both BLEEP and TRIPLEX). The hidden size and dropout rate are set to 512 and 0.1, respectively. For TRIPLEX, we use 4 layers, 8 attention heads, and 25 neighboring spots. For BLEEP, the temperature in the contrastive loss is set to 1, and the number of retrieved spots is 50.

#### Weakly Supervised Classification

Following previous studies [9, 66], we use AdamW as the optimizer with a learning rate of 3 × 10^*−*4^ and weight decay of 1 × 10^*−*5^. A cosine learning rate scheduler is also applied.

## 5 Acknowledgement

We thank Bo Yu for his assistance in curating the gene symbol lists. This work was supported by NSF IIS Div Of Information & Intelligent Systems 2403317 and Amazon research.

## A Appendix

### A.1 Experimental Setup

#### Gene Expression Prediction

We use HEST-Bench^6^ to evaluate STPath on the gene expression prediction task. This benchmark comprises 10 cancer datasets from different organs and provides measured gene expression levels. The original evaluation setup involves assessing performance by averaging the Pearson correlation of the top 50 highly variable genes (HVGs) across multiple folds, with splits performed at the patient level. To prevent label leakage, all HEST-Bench samples are excluded from our pretraining datasets.

In our experiments, instead of using a cross-validation setup, we compare the average performance across all samples on top 50 HVGs without additional fine-tuning. Furthermore, we incorporate adjusted mutual information (AMI) as an additional evaluation metric. This ensures a more realistic evaluation as it reflects the models ability to generalize across diverse datasets, especially in scenarios where downstream training datasets are often unavailable. Error bars represent 95% confidence intervals with nonparametric bootstrapping using 100 bootstrap replicates.

#### Spot Imputation

We adjust the percentage of spots with annotated gene expressions to evaluate STPath’s ability to utilize contextual information. The prompted spots are initially selected from the leftmost spot (i.e., the spot with the minimum x-axis value) and gradually expanded to their nearest neighbors according to the specified percentage. The model’s performance with prompted spots is evaluated solely based on the masked regions. We still apply HEST-Bench and follow the setup of the gene expression prediction task.

#### Spatial Clustering

We use five WSI-ST datasets with pathology annotations: 12 human brain samples with layer annotations [42], 14 GSE213688 samples with breast cancer subtypes [15], 1 Chen sample with tumor annotations [8], 8 Andersson samples with tumor cell types [4], and 7 Erickson samples with benign and malignant tumor annotations [20]. The goal is to determine whether the predicted gene expression effectively captures relevant pathological structures and aligns with expert annotations.

To achieve this, we first predict the gene expressions for all spots in a given WSI and apply Leiden clustering to assign cluster IDs to each spot. The alignment between computed clusters and ground truth annotations is quantified using adjusted mutual information (AMI) and homogeneity score (Homo).

#### Biomarker Prediction

We select several well-established biomarkers from the top 50 HVGs in the HEST-Bench datasets: GATA3 and MYBPC1 for IDC [26, 11, 22], LCN2 for COAD [33], GPRC5A and SOX2 for LUNG [78, 55, 27], and UBE2C for PAAD [70, 41]. Following the same protocol as the gene expression prediction task, we evaluate each model’s ability to predict the expression levels of these biomarkers and compare the predictions with ground truth values. Performance is reported as the average Pearson correlation coefficient across all slides within each dataset. Error bars represent 95% confidence intervals, estimated using nonparametric bootstrapping with 100 replicates.

For biomarker analysis, we performed differential gene expression testing on the predicted spatial transcriptomics from STPath using the Wilcoxon rank-sum test, with region labels derived from pathologist annotations^7^ (on the same datasets used in the spatial clustering task). All genes were included in the analysis, and a minimum log fold-change threshold of 1 was applied to emphasize genes with strong regional specificity. Results were visualized using a matrix plot^8^, which highlights the top 10 marker genes per tissue region based on log fold-change.

#### Weakly Supervised Classification

Each WSI is segmented into spots with a resolution of 256 × 256 pixels at × 10 magnification using the Trident toolkit^9^ [75, 66]. For downstream slide-level classification, we train an attention-based multi-instance learning (ABMIL) head [31], which takes spot-level features as input and aggregates them into a slide-level embedding.

We incorporate the predicted expression of the following biomarkers from STPath as spatial transcriptomic (ST) features: PIK3CA, TP53, BAP1, PBRM1, KRAS, EGFR, CASP8, BRAF, GATA3, MYBPC1, ARID1A, KEAP1, STK11, and SMAD4, most of which are key targets in gene mutation prediction tasks. These ST features are concatenated with visual features for each spot, enriching the representation and providing additional spatial context for the WSI.

AUC is used as the evaluation metric for gene mutation prediction, while the concordance index (c-index) is applied for survival prediction. The best model checkpoint is selected based on the lowest validation loss. All reported results are averaged over three runs with different random seeds to ensure robustness.

#### Unsupervised Gene Mutation Prediction

We use STPath to infer the expression of the target biomarker for each spot. These predicted expressions are aggregated by computing the variance across all spots within a WSI, which serves as the mutation prediction score for that slide. No additional fine-tuning is performed. Evaluation is conducted on all slides in each dataset using this unsupervised prediction approach.

### A.2 Dataset Details

#### Pretraining Dataset

We curate a large-scale collection of H&E-stained histology images annotated with spatial transcriptomics by combining HEST-1K [32] and STImage-1k4m [8]. Since these two datasets include overlapping slides from identical publication sources, we remove any samples in STImage-1k4m that originate from the same sources as those in HEST-1K. The resulting dataset consists of 986 individual slides representing over 17 organs, with major representations from the breast, skin, brain, heart, bowel, lung, prostate, liver, pancreas, kidney, mouth, ovary, uterus, stomach, colon, and eye. The dataset spans four spatial sequencing protocols: Visium HD, Xenium, Visium, and Spatial Transcriptomics. The pretraining dataset includes 38,984 gene symbols, encompassing both reference genes and their isoforms. The overall statistics can be found in Table 1.

**Table 1:**
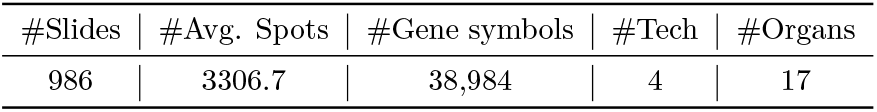
The dataset statistics of the pretrained dataset.

#### Datasets with Expert Pathology Annotations

We study five datasets with pathologistprovided annotations, including both domain-specific and disease-specific labels. These annotations are provided at the spot level, representing localized tissue characteristics such as anatomical regions (e.g., brain layers) or pathological states (e.g., tumor subtypes). The detailed statistics can be found in Table 2.

**Table 2:**
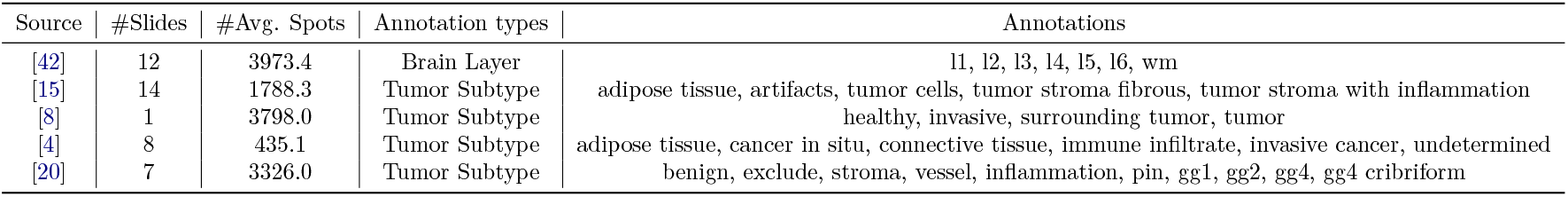
Statistics of the datasets used for spatial segmentation.

In the Erickson dataset [20], labels “gg” refer to the International Society of Urological Pathology (ISUP) Gleason Grade Groups, which are used to grade prostate cancer severity, while “pin” denotes prostatic intraepithelial neoplasia, a precancerous lesion.

#### Slide-level Prediction

The details of all datasets used for gene mutation and survival prediction are summarized in Table 3. The sources for downloading the WSIs are listed in the table, and the corresponding labels are available through the recently released Patho-Bench^10^. For the unsupervised gene mutation prediction task, the entire dataset is used for evaluation. For the weakly supervised prediction tasks, each dataset is randomly split into training, validation, and test sets following a 6/2/2 split at the subject level.

**Table 3:**
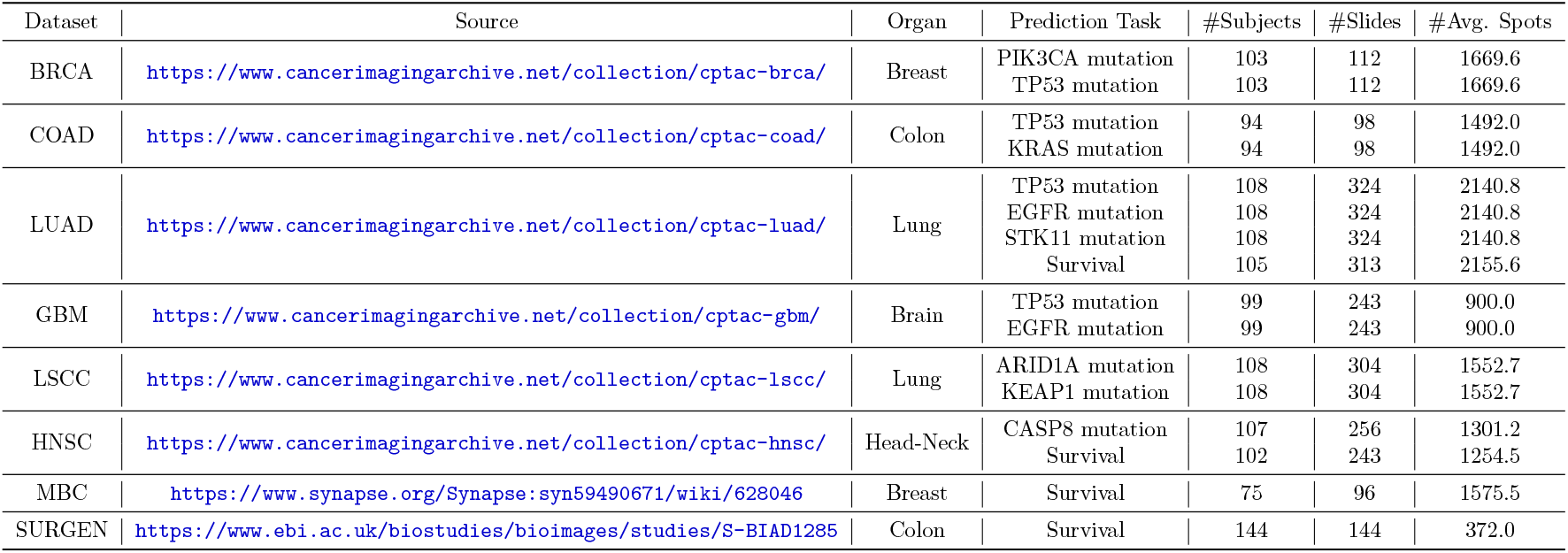
An overview of the datasets used for gene mutation and survival prediction tasks.

### A.3 Full Results of In-context Prediction

Unlike previous studies that focus solely on learning a mapping from images to gene expressions, the generative learning paradigm enables in-context prediction, allowing the model to utilize a few prompted spots’ or genes’ expressions for more informed predictions. In this section, we evaluate the model’s performance in predicting masked spots’ gene expressions based on the expressions of the remaining spots. The ratio of masked spots varies within the range of [5%, 10%, 20%, 30%, 40%, 50%], while a prompted rate of 0% represents the task of predicting expressions for all spots. To ensure a challenging and meaningful prediction setup, masking is applied over continuous regions of spots. The results are presented in Table 4.

**Table 4:** Results of gene expression prediction over the masked spots.

| Prompt% | Metric | IDC | PRAD | PAAD | SKCM | COAD | READ | CCRCC | HCC | LUNG | LYMPH | Average |
| --- | --- | --- | --- | --- | --- | --- | --- | --- | --- | --- | --- | --- |
| 0% | PCC | 0.751<br>±.002 | 0.298<br>±.004 | 0.471<br>±.008 | 0.602<br>±.012 | 0.521<br>±.004 | 0.300<br>±.007 | 0.035<br>±.003 | 0.104<br>±.007 | 0.559<br>±.006 | 0.070<br>±.004 | 0.371 |
|  | AMI | 0.651<br>±.003 | 0.270<br>±.002 | 0.635<br>±.007 | 0.569<br>±.014 | 0.644<br>±.004 | 0.332<br>±.007 | 0.650<br>±.003 | 0.374<br>±.007 | 0.560<br>±.009 | 0.575<br>±.004 | 0.526 |
| 5% | PCC | 0.751<br>±.002 | 0.386<br>±.004 | 0.479<br>±.009 | 0.584<br>±.014 | 0.529<br>±.004 | 0.366<br>±.006 | 0.292<br>±.003 | 0.179<br>±.007 | 0.564<br>±.006 | 0.228<br>±.005 | 0.435 |
|  | AMI | 0.645<br>±.003 | 0.319<br>±.002 | 0.645<br>±.007 | 0.578<br>±.013 | 0.641<br>±.006 | 0.334<br>±.007 | 0.640<br>±.003 | 0.413<br>±.008 | 0.544<br>±.008 | 0.619<br>±.005 | 0.538 |
| 10% | PCC | 0.754<br>±.001 | 0.426<br>±.005 | 0.485<br>±.008 | 0.578<br>±.014 | 0.540<br>±.005 | 0.372<br>±.006 | 0.340<br>±.003 | 0.182<br>±.007 | 0.572<br>±.007 | 0.250<br>±.005 | 0.450 |
|  | AMI | 0.654<br>±.004 | 0.299<br>±.002 | 0.652<br>±.008 | 0.548<br>±.013 | 0.649<br>±.004 | 0.336<br>±.008 | 0.652<br>±.002 | 0.394<br>±.008 | 0.569<br>±.010 | 0.609<br>±.004 | 0.536 |
| 20% | PCC | 0.759<br>±.002 | 0.439<br>±.004 | 0.490<br>±.009 | 0.588<br>±.014 | 0.557<br>±.005 | 0.374<br>±.007 | 0.361<br>±.004 | 0.193<br>±.011 | 0.584<br>±.007 | 0.293<br>±.006 | 0.463 |
|  | AMI | 0.669<br>±.003 | 0.333<br>±.003 | 0.639<br>±.009 | 0.554<br>±.013 | 0.642<br>±.005 | 0.346<br>±.009 | 0.671<br>±.004 | 0.416<br>±.012 | 0.577<br>±.010 | 0.619<br>±.004 | 0.546 |
| 30% | PCC | 0.762<br>±.002 | 0.433<br>±.005 | 0.496<br>±.010 | 0.602<br>±.012 | 0.577<br>±.005 | 0.369<br>±.007 | 0.375<br>±.003 | 0.189<br>±.007 | 0.589<br>±.007 | 0.316<br>±.006 | 0.470 |
|  | AMI | 0.685<br>±.004 | 0.304<br>±.003 | 0.662<br>±.010 | 0.574<br>±.016 | 0.627<br>±.006 | 0.345<br>±.008 | 0.646<br>±.003 | 0.401<br>±.009 | 0.596<br>±.012 | 0.630<br>±.005 | 0.547 |
| 40% | PCC | 0.766<br>±.004 | 0.446<br>±.003 | 0.489<br>±.010 | 0.613<br>±.016 | 0.593<br>±.006 | 0.349<br>±.008 | 0.387<br>±.003 | 0.187<br>±.009 | 0.594<br>±.012 | 0.334<br>±.005 | 0.476 |
|  | AMI | 0.666<br>±.004 | 0.327<br>±.003 | 0.650<br>±.010 | 0.594<br>±.017 | 0.636<br>±.006 | 0.331<br>±.010 | 0.688<br>±.003 | 0.413<br>±.012 | 0.609<br>±.013 | 0.631<br>±.005 | 0.554 |
| 50% | PCC | 0.771<br>±.002 | 0.477<br>±.008 | 0.483<br>±.009 | 0.620<br>±.015 | 0.603<br>±.006 | 0.339<br>±.011 | 0.399<br>±.005 | 0.182<br>±.009 | 0.603<br>±.008 | 0.341<br>±.007 | 0.481 |
|  | AMI | 0.642<br>±.002 | 0.311<br>±.008 | 0.661<br>±.009 | 0.598<br>±.015 | 0.626<br>±.006 | 0.351<br>±.011 | 0.707<br>±.005 | 0.429<br>±.009 | 0.612<br>±.008 | 0.643<br>±.007 | 0.558 |

<sup>10</sup>https://huggingface.co/datasets/MahmoodLab/Patho-Bench

## Footnotes

1 We exclude BLEEP since its pretraining process is not stable.

2 https://github.com/mahmoodlab/UNI

3 https://github.com/prov-gigapath/prov-gigapath

4 https://github.com/bowang-lab/BLEEP/tree/main

5 https://github.com/NEXGEM/TRIPLEX/tree/main

6 https://huggingface.co/datasets/MahmoodLab/hest-bench/tree/main

7 https://scanpy.readthedocs.io/en/stable/generated/scanpy.tl.rank_genes_groups.html

8 https://scanpy.readthedocs.io/en/stable/api/generated/scanpy.pl.rank_genes_groups_matrixplot.html

9 https://github.com/mahmoodlab/TRIDENT

10 https://huggingface.co/datasets/MahmoodLab/Patho-Bench

## References

[1] Sarmad Al-Marsoummi, Emilie Vomhof-DeKrey, and Marc D Basson. Schlafen12 reduces the aggressiveness of triple negative breast cancer through post-transcriptional regulation of zeb1 that drives stem cell differentiation. Cellular physiology and biochemistry: international journal of experimental cellular physiology, biochemistry, and pharmacology, 53(6):999, 2019.

[2] Farin F Amersi, Alicia M Terando, Yasufumi Goto, Richard A Scolyer, John F Thompson, Andy N Tran, Mark B Faries, Donald L Morton, and Dave SB Hoon. Activation of ccr9/ccl25 in cutaneous melanoma mediates preferential metastasis to the small intestine. Clinical cancer research, 14(3):638–645, 2008.

[3] Javeria Amin, Muhammad Sharif, Mussarat Yasmin, and Steven Lawrence Fernandes. A distinctive approach in brain tumor detection and classification using mri. Pattern Recognition Letters, 139:118–127, 2020.

[4] Alma Andersson, Ludvig Larsson, Linnea Stenbeck, Fredrik Salmén, Anna Ehinger, Sunny Z Wu, Ghamdan Al-Eryani, Daniel Roden, Alex Swarbrick, Åke Borg, et al. Spatial deconvolution of her2-positive breast cancer delineates tumor-associated cell type interactions. Nature communications, 12(1):6012, 2021.

[5] Kaustav Bera, Kurt A Schalper, David L Rimm, Vamsidhar Velcheti, and Anant Madabhushi. Artificial intelligence in digital pathologynew tools for diagnosis and precision oncology. Nature reviews Clinical oncology, 16(11):703–715, 2019.

[6] Erik N Bergstrom, Ammal Abbasi, Marcos Díaz-Gay, Loïck Galland, Sylvain Ladoire, Scott M Lippman, and Ludmil B Alexandrov. Deep learning artificial intelligence predicts homologous recombination deficiency and platinum response from histologic slides. Journal of Clinical Oncology, 42(30):3550–3560, 2024.

[7] Robert W Carlson, D Craig Allred, Benjamin O Anderson, Harold J Burstein, W Bradford Carter, Stephen B Edge, John K Erban, William B Farrar, Andres Forero, Sharon Hermes Giordano, et al. Invasive breast cancer. Journal of the National Comprehensive Cancer Network, 9(2):136–222, 2011.

[8] Jiawen Chen, Muqing Zhou, Wenrong Wu, Jinwei Zhang, Yun Li, and Didong Li. Stimage-1k4m: A histopathology image-gene expression dataset for spatial transcriptomics. arXiv preprint arXiv:2406.06393, 2024.

[9] Richard J Chen, Tong Ding, Ming Y Lu, Drew FK Williamson, Guillaume Jaume, Bowen Chen, Andrew Zhang, Daniel Shao, Andrew H Song, Muhammad Shaban, et al. Towards a general-purpose foundation model for computational pathology. Nature Medicine, 2024.

[10] Wei-Ting Chen, Ashley Lu, Katleen Craessaerts, Benjamin Pavie, Carlo Sala Frigerio, Nikky Corthout, Xiaoyan Qian, Jana Laláková, Malte Kühnemund, Iryna Voytyuk, et al. Spatial transcriptomics and in situ sequencing to study alzheimers disease. Cell, 182(4):976–991, 2020.

[11] Xiuwen Chen, Weilin Zhao, Yugang Huang, Senyuan Luo, Xianbin Tang, and Qiong Yi. Association of gata3 expression in triple-positive breast cancer with overall survival and immune cell infiltration. Scientific Reports, 14(1):17795, 2024.

[12] Youngmin Chung, Ji Hun Ha, Kyeong Chan Im, and Joo Sang Lee. Accurate spatial gene expression prediction by integrating multi-resolution features. In Proceedings of the IEEE/CVF Conference on Computer Vision and Pattern Recognition, pages 11591–11600, 2024.

[13] GTEx Consortium, Kristin G Ardlie, David S Deluca, Ayellet V Segrè, Timothy J Sullivan, Taylor R Young, Ellen T Gelfand, Casandra A Trowbridge, Julian B Maller, Taru Tukiainen, et al. The genotype-tissue expression (gtex) pilot analysis: multitissue gene regulation in humans. Science, 348(6235):648–660, 2015.

[14] National Cancer Institute Clinical Proteomic Tumor Analysis Consortium et al. The clinical proteomic tumor analysis consortium glioblastoma multiforme collection (cptac-gbm)(version 16)[data set]. the cancer imaging archive. 2018.

[15] Angèle Coutant, Vincent Cockenpot, Lauriane Muller, Cyril Degletagne, Roxane Pommier, Laurie Tonon, Maude Ardin, Marie-Cécile Michallet, Christophe Caux, Marie Laurent, et al. Spatial transcriptomics reveal pitfalls and opportunities for the detection of rare high-plasticity breast cancer subtypes. Laboratory investigation, 103(12):100258, 2023.

[16] Charles Frederick Albert Culling, RT Allison, and WT Barr. Cellular pathology technique. Elsevier, 2014.

[17] Nathan J Edwards, Mauricio Oberti, Ratna R Thangudu, Shuang Cai, Peter B McGarvey, Shine Jacob, Subha Madhavan, and Karen A Ketchum. The cptac data portal: a resource for cancer proteomics research. Journal of proteome research, 14(6):2707–2713, 2015.

[18] Chee-Huat Linus Eng, Michael Lawson, Qian Zhu, Ruben Dries, Noushin Koulena, Yodai Takei, Jina Yun, Christopher Cronin, Christoph Karp, Guo-Cheng Yuan, et al. Transcriptome-scale super-resolved imaging in tissues by rna seqfish+. Nature, 568(7751):235–239, 2019.

[19] Camilla Engblom and Joakim Lundeberg. Putting cancer immunotherapy into spatial context in the clinic. Nature Biotechnology, 43(4):471–476, 2025.

[20] Andrew Erickson, Mengxiao He, Emelie Berglund, Maja Marklund, Reza Mirzazadeh, Niklas Schultz, Linda Kvastad, Alma Andersson, Ludvig Bergenstråhle, Joseph Bergenstråhle, et al. Spatially resolved clonal copy number alterations in benign and malignant tissue. Nature, 608(7922):360–367, 2022.

[21] Xiaofeng Gao, Huitong Bu, Xuzheng Gao, Ying Wang, Long Wang, and Zhenwang Zhang. Pancancer analysis: Spag5 is an immunological and prognostic biomarker for multiple cancers. The FASEB Journal, 37(10):e23159, 2023.

[22] Janelle Geist and Aikaterini Kontrogianni-Konstantopoulos. Mybpc1, an emerging myopathic gene: what we know and what we need to learn. Frontiers in Physiology, 7:410, 2016.

[23] Thomas Hayes, Roshan Rao, Halil Akin, Nicholas J Sofroniew, Deniz Oktay, Zeming Lin, Robert Verkuil, Vincent Q Tran, Jonathan Deaton, Marius Wiggert, et al. Simulating 500 million years of evolution with a language model. Science, page eads0018, 2025.

[24] Bryan He, Ludvig Bergenstråhle, Linnea Stenbeck, Abubakar Abid, Alma Andersson, Åke Borg, Jonas Maaskola, Joakim Lundeberg, and James Zou. Integrating spatial gene expression and breast tumour morphology via deep learning. Nature biomedical engineering, 4(8):827–834, 2020.

[25] Céline N Heinz, Amelie Echle, Sebastian Foersch, Andrey Bychkov, and Jakob Nikolas Kather. The future of artificial intelligence in digital pathology–results of a survey across stakeholder groups. Histopathology, 80(7):1121–1127, 2022.

[26] I-Cheng Ho, Tzong-Shyuan Tai, and Sung-Yun Pai. Gata3 and the t-cell lineage: essential functions before and after t-helper-2-cell differentiation. Nature reviews immunology, 9(2):125–135, 2009.

[27] Zachariah E Holmes, Desmond J Hamilton, Taeyoung Hwang, Nicholas V Parsonnet, John L Rinn, Deborah S Wuttke, and Robert T Batey. The sox2 transcription factor binds rna. Nature communications, 11(1):1805, 2020.

[28] Tinglin Huang, Zhenqiao Song, Rex Ying, and Wengong Jin. Protein-nucleic acid complex modeling with frame averaging transformer. In Advances in Neural Information Processing Systems, 2024.

[29] Nguyen Tho Huu, Hideki Yoshida, and Masamitsu Yamaguchi. Tumor suppressor gene oscp 1/nor 1 regulates apoptosis, proliferation, differentiation, and ros generation during eye development of drosophila melanogaster. The FEBS Journal, 282(24):4727–4746, 2015.

[30] William L Hwang, Karthik A Jagadeesh, Jimmy A Guo, Hannah I Hoffman, Payman Yadollahpour, Jason W Reeves, Rahul Mohan, Eugene Drokhlyansky, Nicholas Van Wittenberghe, Orr Ashenberg, et al. Single-nucleus and spatial transcriptome profiling of pancreatic cancer identifies multicellular dynamics associated with neoadjuvant treatment. Nature genetics, 54(8):1178–1191, 2022.

[31] Maximilian Ilse, Jakub Tomczak, and Max Welling. Attention-based deep multiple instance learning. In International conference on machine learning, pages 2127–2136. PMLR, 2018.

[32] Guillaume Jaume, Paul Doucet, Andrew H. Song, Ming Y. Lu, Cristina Almagro-Perez, Sophia J. Wagner, Anurag J. Vaidya, Richard J. Chen, Drew F. K. Williamson, Ahrong Kim, and Faisal Mahmood. Hest-1k: A dataset for spatial transcriptomics and histology image analysis. In Advances in Neural Information Processing Systems, December 2024.

[33] Byung-Kwon Jung and Kwon-Yul Ryu. Lipocalin-2: a therapeutic target to overcome neurodegenerative diseases by regulating reactive astrogliosis. Experimental & Molecular Medicine, 55(10):2138–2146, 2023.

[34] Joshua S Kaminker, Yan Zhang, Colin Watanabe, and Zemin Zhang. Canpredict: a computational tool for predicting cancer-associated missense mutations. Nucleic acids research, 35(suppl_2):W595–W598, 2007.

[35] Jacob Devlin Ming-Wei Chang Kenton and Lee Kristina Toutanova. Bert: Pre-training of deep bidirectional transformers for language understanding. In Proceedings of naacL-HLT, volume 1, page 2. Minneapolis, Minnesota, 2019.

[36] John M Kirkwood, M Hunt Strawderman, Marc S Ernstoff, Thomas J Smith, Ernest C Borden, and Ronald H Blum. Interferon alfa-2b adjuvant therapy of high-risk resected cutaneous melanoma: the eastern cooperative oncology group trial est 1684. Journal of clinical oncology, 14(1):7–17, 1996.

[37] Alona Levy-Jurgenson, Xavier Tekpli, Vessela N Kristensen, and Zohar Yakhini. Spatial transcriptomics inferred from pathology whole-slide images links tumor heterogeneity to survival in breast and lung cancer. Scientific reports, 10(1):18802, 2020.

[38] Zhaoting Li, Fanyi Mo, Yixin Wang, Wen Li, Yu Chen, Jun Liu, Ting-Jing Chen-Mayfield, and Quanyin Hu. Enhancing gasdermin-induced tumor pyroptosis through preventing escrt-dependent cell membrane repair augments antitumor immune response. Nature Communications, 13(1):6321, 2022.

[39] Tianyu Liu, Tinglin Huang, Rex Ying, and Hongyu Zhao. spemo: Exploring the capacity of foundation models for analyzing spatial multi-omic data. bioRxiv, pages 2025–01, 2025.

[40] Ming Y Lu, Bowen Chen, Drew FK Williamson, Richard J Chen, Ivy Liang, Tong Ding, Guillaume Jaume, Igor Odintsov, Long Phi Le, Georg Gerber, et al. A visual-language foundation model for computational pathology. Nature Medicine, 30(3):863–874, 2024.

[41] Sijia Ma, Qian Chen, Xu Li, Jing Fu, and Le Zhao. Ube2c serves as a prognosis biomarker of uterine corpus endometrial carcinoma via promoting tumor migration and invasion. Scientific Reports, 13(1):16899, 2023.

[42] Kristen R Maynard, Leonardo Collado-Torres, Lukas M Weber, Cedric Uytingco, Brianna K Barry, Stephen R Williams, Joseph L Catallini, Matthew N Tran, Zachary Besich, Madhavi Tippani, et al. Transcriptome-scale spatial gene expression in the human dorsolateral prefrontal cortex. Nature neuroscience, 24(3):425–436, 2021.

[43] Mayur D Mody, James W Rocco, Sue S Yom, Robert I Haddad, and Nabil F Saba. Head and neck cancer. The Lancet, 398(10318):2289–2299, 2021.

[44] Jeffrey R Moffitt, Dhananjay Bambah-Mukku, Stephen W Eichhorn, Eric Vaughn, Karthik Shekhar, Julio D Perez, Nimrod D Rubinstein, Junjie Hao, Aviv Regev, Catherine Dulac, et al. Molecular, spatial, and functional single-cell profiling of the hypothalamic preoptic region. Science, 362(6416):eaau5324, 2018.

[45] Sébastien Molière, Massimo Lodi, Suzanne Leblanc, Anne Gressel, Carole Mathelin, Fabien Alpy, Marie-Pierre Chenard, and Catherine Tomasetto. Mmp-11 expression in early luminal breast cancer: associations with clinical, mri, pathological characteristics, and disease-free survival. BMC cancer, 24(1):295, 2024.

[46] David J Myers and Jason M Wallen. Lung adenocarcinoma. In StatPearls [Internet]. StatPearls Publishing, 2023.

[47] Craig Myles, In Hwa Um, Craig Marshall, David Harris-Birtill, and David J Harrison. Surgen: 1020 h&e-stained whole slide images with survival and genetic markers. arXiv preprint 2502.04946, 2025.

[48] Omri Puny, Matan Atzmon, Heli Ben-Hamu, Ishan Misra, Aditya Grover, Edward J Smith, and Yaron Lipman. Frame averaging for invariant and equivariant network design. arXiv preprint 2110.03336, 2021.

[49] Hui Qu, Mu Zhou, Zhennan Yan, He Wang, Vinod K Rustgi, Shaoting Zhang, Olivier Gevaert, and Dimitris N Metaxas. Genetic mutation and biological pathway prediction based on whole slide images in breast carcinoma using deep learning. NPJ precision oncology, 5(1):87, 2021.

[50] Zefang Ren, Aesun Shin, Qiuyin Cai, Xiao-Ou Shu, Yu-Tang Gao, and Wei Zheng. Igfbp3 mrna expression in benign and malignant breast tumors. Breast Cancer Research, 9:1–9, 2007.

[51] Boris Reva, Yevgeniy Antipin, and Chris Sander. Predicting the functional impact of protein mutations: application to cancer genomics. Nucleic acids research, 39(17):e118–e118, 2011.

[52] Samuel G Rodriques, Robert R Stickels, Aleksandrina Goeva, Carly A Martin, Evan Murray, Charles R Vanderburg, Joshua Welch, Linlin M Chen, Fei Chen, and Evan Z Macosko. Slide-seq: A scalable technology for measuring genome-wide expression at high spatial resolution. Science, 363(6434):1463–1467, 2019.

[53] I Salama, PS Malone, F Mihaimeed, and i JL Jones. A review of the s100 proteins in cancer. European Journal of Surgical Oncology (EJSO), 34(4):357–364, 2008.

[54] Anna C Schaar, Alejandro Tejada-Lapuerta, Giovanni Palla, Robert Gutgesell, Lennard Halle, Mariia Minaeva, Larsen Vornholz, Leander Dony, Francesca Drummer, Mojtaba Bahrami, et al. Nicheformer: a foundation model for single-cell and spatial omics. bioRxiv, pages 2024–04, 2024.

[55] Thorsten Schaefer and Claudia Lengerke. Sox2 protein biochemistry in stemness, reprogramming, and cancer: the pi3k/akt/sox2 axis and beyond. Oncogene, 39(2):278–292, 2020.

[56] Stuart J Schnitt. Classification and prognosis of invasive breast cancer: from morphology to molecular taxonomy. Modern pathology, 23:S60–S64, 2010.

[57] Zhuchen Shao, Yang Chen, Hao Bian, Jian Zhang, Guojun Liu, and Yongbing Zhang. Hvtsurv: Hierarchical vision transformer for patient-level survival prediction from whole slide image. In Proceedings of the AAAI Conference on Artificial Intelligence, volume 37, pages 2209–2217, 2023.

[58] Andrew H Song, Guillaume Jaume, Drew FK Williamson, Ming Y Lu, Anurag Vaidya, Tiffany R Miller, and Faisal Mahmood. Artificial intelligence for digital and computational pathology. Nature Reviews Bioengineering, 1(12):930–949, 2023.

[59] Patrik L Ståhl, Fredrik Salmén, Sanja Vickovic, Anna Lundmark, José Fernández Navarro, Jens Magnusson, Stefania Giacomello, Michaela Asp, Jakub O Westholm, Mikael Huss, et al. Visualization and analysis of gene expression in tissue sections by spatial transcriptomics. Science, 353(6294):78–82, 2016.

[60] Jan M Suski, Nalin Ratnayeke, Marcin Braun, Tian Zhang, Vladislav Strmiska, Wojciech Michowski, Geylani Can, Antoine Simoneau, Konrad Snioch, Mikolaj Cup, et al. Cdc7-independent g1/s transition revealed by targeted protein degradation. Nature, 605(7909):357–365, 2022.

[61] Ratna Rajesh Thangudu, Paul A Rudnick, Michael Holck, Deepak Singhal, Michael J MacCoss, Nathan J Edwards, Karen A Ketchum, Christopher R Kinsinger, Erika Kim, and Anand Basu. Abstract lb-242: Proteomic data commons: A resource for proteogenomic analysis. Cancer Research, 80(16_Supplement): LB–242, 2020.

[62] Christina V Theodoris, Ling Xiao, Anant Chopra, Mark D Chaffin, Zeina R Al Sayed, Matthew C Hill, Helene Mantineo, Elizabeth M Brydon, Zexian Zeng, X Shirley Liu, et al. Transfer learning enables predictions in network biology. Nature, 618(7965):616–624, 2023.

[63] Baldwin H Tom, Lynne P Rutzky, Milda M Jakstys, Ryoichi Oyasu, Celia I Kaye, and Barry D Kahan. Human colonic adenocarcinoma cells: I. establishment and description of a new line. In vitro, 12:180–191, 1976.

[64] Katarzyna Tomczak, Patrycja Czerwińska, and Maciej Wiznerowicz. Review the cancer genome atlas (tcga): an immeasurable source of knowledge. Contemporary Oncology/Współczesna Onkologia, 2015(1):68–77, 2015.

[65] Kaja Urbańska, Justyna Sokołowska, Maciej Szmidt, and Paweł Sysa. Glioblastoma multiforme–an overview. Contemporary Oncology/Współczesna Onkologia, 18(5):307–312, 2014.

[66] Anurag Vaidya, Andrew Zhang, Guillaume Jaume, Andrew H Song, Tong Ding, Sophia J Wagner, Ming Y Lu, Paul Doucet, Harry Robertson, Cristina Almagro-Perez, et al. Molecular-driven foundation model for oncologic pathology. arXiv preprint 2501.16652, 2025.

[67] Ashish Vaswani, Noam Shazeer, Niki Parmar, Jakob Uszkoreit, Llion Jones, Aidan N Gomez, łukasz Kaiser, and Illia Polosukhin. Attention is all you need. Advances in neural information processing systems, 30, 2017.

[68] Rupert Allan Willis. Pathology of tumours. 1948.

[69] Ellery Wulczyn, David F Steiner, Melissa Moran, Markus Plass, Robert Reihs, Fraser Tan, Isabelle Flament-Auvigne, Trissia Brown, Peter Regitnig, Po-Hsuan Cameron Chen, et al. Interpretable survival prediction for colorectal cancer using deep learning. NPJ digital medicine, 4(1):71, 2021.

[70] Cheng Xiang and Hai-chao Yan. Ubiquitin conjugating enzyme e2 c (ube2c) may play a dual role involved in the progression of thyroid carcinoma. Cell Death Discovery, 8(1):130, 2022.

[71] Yi Xiao and Dihua Yu. Tumor microenvironment as a therapeutic target in cancer. Pharmacology & therapeutics, 221:107753, 2021.

[72] Ronald Xie, Kuan Pang, Sai Chung, Catia Perciani, Sonya MacParland, Bo Wang, and Gary Bader. Spatially resolved gene expression prediction from histology images via bi-modal contrastive learning. Advances in Neural Information Processing Systems, 36, 2024.

[73] Hanwen Xu, Naoto Usuyama, Jaspreet Bagga, Sheng Zhang, Rajesh Rao, Tristan Naumann, Cliff Wong, Zelalem Gero, Javier González, Yu Gu, Yanbo Xu, Mu Wei, Wenhui Wang, Shuming Ma, Furu Wei, Jianwei Yang, Chunyuan Li, Jianfeng Gao, Jaylen Rosemon, Tucker Bower, Soohee Lee, Roshanthi Weerasinghe, Bill J. Wright, Ari Robicsek, Brian Piening, Carlo Bifulco, Sheng Wang, and Hoifung Poon. A whole-slide foundation model for digital pathology from real-world data. Nature, 2024.

[74] Jingrun Yang, Yang Xu, Kuixia Xie, Ling Gao, Wenying Zhong, and Xinhua Liu. Cebpb is associated with active tumor immune environment and favorable prognosis of metastatic skin cutaneous melanoma. Frontiers in Immunology, 13:991797, 2022.

[75] Andrew Zhang, Guillaume Jaume, Anurag Vaidya, Tong Ding, and Faisal Mahmood. Accelerating data processing and benchmarking of ai models for pathology. arXiv preprint 2502.06750, 2025.

[76] Linlin Zhang, Dongsheng Chen, Dongli Song, Xiaoxia Liu, Yanan Zhang, Xun Xu, and Xiangdong Wang. Clinical and translational values of spatial transcriptomics. Signal Transduction and Targeted Therapy, 7(1):111, 2022.

[77] Xu Zhang, Shuai Huang, Junchao Guo, Li Zhou, Lei You, Taiping Zhang, and Yupei Zhao. Insights into the distinct roles of mmp-11 in tumor biology and future therapeutics. International journal of oncology, 48(5):1783–1793, 2016.

[78] H Zhou, AG Telonis, Y Jing, NL Xia, L Biederman, M Jimbo, F Blanco, E Londin, JR Brody, and I Rigoutsos. Gprc5a is a potential oncogene in pancreatic ductal adenocarcinoma cells that is upregulated by gemcitabine with help from hur. Cell death & disease, 7(7):e2294–e2294, 2016.

[79] Xiaoli Zhou, Lizhou Jia, Yangyang Sun, Lingyun Xu, Xudong Wang, and Qi Tang. Sperm-associated antigen 5 is a potential biomarker for poor prognosis in breast cancer. Oncology Letters, 17(1):1146–1152, 2019.

[80] Eric Zimmermann, Eugene Vorontsov, Julian Viret, Adam Casson, Michal Zelechowski, George Shaikovski, Neil Tenenholtz, James Hall, Thomas Fuchs, Nicolo Fusi, Siqi Liu, and Kristen Severson. Virchow2: Scaling self-supervised mixed magnification models in pathology. arXiv preprint 2408.00738, 2024.

